# MetaMAG Explorer: A Database-Augmenting Pipeline for Genome-Resolved Metagenomics and Enhanced Microbial Classification

**DOI:** 10.64898/2026.04.27.721001

**Authors:** Maria Altaf Satti, Zexi Cai

## Abstract

Accurate taxonomic classification in metagenomic studies remains challenging because reference databases are often static and incomplete, limiting our understanding of microbial diversity, especially in habitats that are not well represented. We introduce MetaMAG Explorer, a complete and modular pipeline designed to fill this gap with its unique database augmentation framework. Together with end-to-end features like read preprocessing, assembly, binning, and annotation, MetaMAG also presents an automated method for finding new metagenome-assembled genomes (MAGs), confirming their uniqueness by dereplication against curated repositories, and dynamically adding them to classification databases that are compatible with Kraken2. Additionally, MetaMAG makes it easier to understand data by automatically creating high-quality figures that are ready for publication, allowing results to be quickly included in scientific papers. Evaluated across human, plant, and rumen datasets, MetaMAG recovered 233 MAGs, including 121 high-quality genomes, of which 48 (20%) were novel. Database augmentation increased Kraken2 classification rates and reassigned millions of previously misclassified reads. Beyond the gain in read classification, the database augmentation revealed ecologically important taxa that are consistently present in all samples but previously undetected. By enabling iterative database growth driven by the novel MAGs, MetaMAG offers a scalable, highly reproducible, and extensible solution for truly genome-resolved metagenomics, advancing both microbial discovery and taxonomic classification accuracy.

## Introduction

Metagenomics revolutionized microbial ecology by enabling the analysis of complex microbial communities directly from environmental samples. The most important aspect of its feature is the reconstruction of metagenome-assembled genomes (MAGs), in which it is feasible to obtain genomic-level information without cultivation. MAGs have further broadened our understanding of the microbial tree of life [1], revealed novel taxa from human [2], environmental [3], and host-microbial microbiomes [4], and contributed to the growing inventory of uncultivated microbial lineages [5]. Despite these advances, a major obstacle remains: taxonomic classification of metagenomic reads still relies on incomplete reference databases that are heavily biased toward cultured organisms and rarely updated with project-specific data. Accordingly, a significant proportion of metagenomic data, sometimes over 50%, remains unclassifiable [6, 7], particularly in environments such as the bovine rumen [5], plant rhizosphere [8], or non-Western gut microbiota [2, 3]. Tools like Kraken2 [9], although fast and efficient, are still limited by the incompleteness of the reference databases they rely on.

The issue arises in complex environments like the rumen microbiome, where limited databases make it harder to classify microbes accurately. Smith et al. (2022) have established that the precision of rumen metagenomic data classification is significantly dependent on the reference database. Using simulated data, they found that standard databases such as NCBI RefSeq assigned fewer than 40% of the reads to the genus level and regularly misassigned species-level reads. Adding carefully chosen isolates from the Hungate collection to the database improved the results. Additionally beneficial were rumen-specific MAGs (RUGs) [10]. These findings emphasize the need for tools that can successfully integrate novel MAGs into reference databases and recover them for improved classification accuracy in underrepresented environments.

Several solutions have tried to address this challenge with added taxonomic infrastructure (e.g., the Genome Taxonomy Database - GTDB) [11] and more comprehensive MAG databases [2, 3]. Still, most tools don’t support automatically adding new MAGs to their classification databases, which limits taxonomic classification and downstream analysis of novel organisms. Existing pipelines such as SqueezeMeta [12] and ATLAS [13] provide useful frameworks for assembly, binning, and annotation, while METABOLIC [14] focuses on functional trait profiling. EasyMetagenome [15], in contrast, creates an easy-to-use interface that integrates read-based and assembly-based analysis with visualization. Although helpful, such pipelines generally lack methodologies for the automatic integration of newly recovered MAGs into reference databases, so a large proportion of reads are left unclassified in complex environments.

To bridge this gap, we developed MetaMAG Explorer, an entirely modular pipeline that goes beyond standard MAG recovery. Although MetaMAG contains all the standard stages of MAG analysis: quality filtering, assembly, binning, dereplication, taxonomic assignment, and functional annotation, its biggest added value is in its module for database augmentation. This pipeline identifies putatively novel MAGs by GTDB-Tk taxonomy assignment [16], dereplicates against curated repositories (including environment-specific libraries), and formats and merges these MAGs automatically into Kraken2-compatible reference databases [9]. This enrichment pipeline is paired with a taxonomy reconciliation system to transform the results of GTDB into NCBI taxdump structures, assign unique identifiers to novel taxa, and update the Kraken2 database in return. The result is a dynamically improving classification framework that improves with each round of analysis, enabling ongoing optimization of microbial reference catalogs.

We tested MetaMAG on metagenomic samples from three representative environments: the human gut [17], the plant rhizosphere [18], and the bovine rumen [19]. For each of these environments, MetaMAG recovered several high-quality MAGs, many of which had no species-level classification in GTDB. Including these MAGs in the classification database gave substantial read-level assignments and allowed for the detection of new microbial clades. Our results indicate that the database expansion strategy of MetaMAG enhances both microbial discovery and downstream taxonomic profiling quality.

## Methods

### Pipeline Architecture and Implementation

MetaMAG Explorer is a computational pipeline for end-to-end metagenome-assembled genome (MAG) analysis with integrated database augmentation capabilities. The core workflow is divided into seven processing stages: (1) data preprocessing, (2) assembly, (3) binning and refinement, (4) novel MAG detection and database augmentation, (5) functional annotation, (6) phylogeny, and (7) abundance estimation. These stages are implemented as fifteen modular components which can be run independently or as part of a fully automated SLURM-managed workflow. Each step records tool versions, parameters, and metadata for full reproducibility.

### Installation and Deployment Framework

MetaMAG Explorer uses Conda to manage software dependencies. Instead of one fully automated installation, the setup works through helper scripts such as setup_tools.py. These scripts allow installation of individual tools for tasks like trimming, assembly, or annotation, as well as entire groups of tools when needed. Reference databases are not included by default, but scripts are provided to download and install them. The same scripts also check that installations are complete and correctly configured. The pipeline is designed for SLURM-managed HPC systems. Updates to tools or databases must be done manually by re-running the appropriate setup commands.

### Core Software Integration and Database Resources

The pipeline integrates major tools for each of the steps involved, such as preprocessing, assembly, binning, refinement, quality control, taxonomic classification, and functional annotation. FastQC, fastp, BWA-mem, MEGAHIT, MetaBAT2, CheckM2, GTDB-Tk, and Kraken2 are some of the tools integrated to facilitate analysis. The pipeline is constructed to process and integrate several reference databases that are required for in-depth MAG analysis. Although these databases are not provided directly with the pipeline, the pipeline offers scripts that can be utilized for downloading and installing them. The important resources are GTDB r226 for taxonomy annotation (over 400,000 bacterial and archaeal genomes), eggNOG v5.0 for orthology mapping (exceeding 5 million orthologous groups), KEGG for pathway reconstruction (more than 500 reference pathways), CAZy for carbohydrate-active enzyme annotation (surpassing 600 enzyme families), and Kraken2 databases for diverse analytical contexts. For rumen-specific studies, scripts are available to integrate reference libraries, including RUG (4,941 genomes), RMGMC (10,373 genomes), and the MGnify Cow Rumen Species Catalogue (2,729 genomes). **Figure S1** illustrates how each pipeline module connects to its associated tools and databases, with dashed arrows indicating specific database dependencies.

### Pipeline Workflow

MetaMAG Explorer was created as a modular metagenomic analysis pipeline that integrates built-in database augmentation features. The pipeline has seven modules: quality control, assembly, binning, MAG quality assessment, novel MAG detection, database augmentation, functional annotation, and abundance profiling. Python and Bash scripting were used to implement the workflow, and SLURM job scheduling was used to control its execution on high-performance computing clusters.

### Phase 1: Data Preprocessing

The preprocessing module starts with FastQC [20] (default settings), which checks read quality and generates detailed reports on sequence quality metrics, adapter contamination, and potential sequencing artifacts. Next, fastp [21] is used to remove adapters, trim low-quality bases (below Q20), cut very short reads (under 50 bp), and filter out low-complexity sequences. Host DNA is then removed using BWA-mem [22], which aligns reads to a provided host genome e.g. (GRCh38 for human [23], ARS-UCD1.2 [24] for bovine, and Lactuca sativa v8 [25] for lettuce). SAMtools [26] (samtools view -f 4), took out reads that didn’t line up with the host and changed them to FASTQ format. FastQC was used to check the quality of the preprocessing again on clean reads, this cleaned set of reads is used for assembly and further analysis.

### Phase 2: Assembly

There are two choices for assembly: single assembly and co-assembly. The user can choose either one method or both. IDBA-UD [27] is used for single-sample assembly, which helps identify genomes that are common and well-covered. MEGAHIT [28] is used for co-assembly, which uses reads from multiple samples to identify genomes that are rare or not as common. The assemblies are then checked with MetaQUAST (v5.2.0) [29] to see if they are full, correct, and free of any errors.

### Phase 3: Binning and Refinement

For binning modules in MetaWRAP [30] were used. That uses three algorithms to do genome binning: MetaBAT2 [31] (which clusters based on tetranucleotide frequency and coverage), MaxBin2 [32] (which uses expectation-maximization), and CONCOCT [33] (which uses composition-based clustering). DAS Tool [34] was used for bin refinement. MAGs were dereplicated using dRep [35] and evaluated for contamination and completeness using CheckM2 [36].

### Phase 4: Novel MAG Detection and Database Augmentation

The classify_wf workflow of GTDB-Tk (v2.3.0) [16] was used for taxonomic assignment of dereplicated MAGs. We flagged MAGs as potentially novel when GTDB-Tk couldn’t assign them to any known species (classification strings ending with “s_”), indicating they had no close relatives in the GTDB reference database at the species level (typically <95% ANI to any reference genome). For rumen samples, novel candidates were subjected to additional dereplication against specialized rumen reference genome collections. Reference datasets were acquired using multi-threaded downloading: RGMGC [37] genomes were downloaded from the ENA browser [38] API using the Python requests library. MGnify [39, 40] cow-rumen genomes were retrieved from EBI FTP servers using recursive web scraping with BeautifulSoup, and the RUG dataset [41] was obtained using wget from DataShare Edinburgh [42]. The downloaded archives were converted to the .fa format, decompressed using gunzip, and extracted using tar.

Using dRep with fastANI [43] as the secondary clustering algorithm, dereplication was carried out with the following thresholds applied: 90% ANI similarity, 10% maximum contamination, 100% strain heterogeneity, and 80% minimum completeness. Project MAGs that remained after dereplication against these comprehensive reference datasets were considered truly novel and project-specific genomes. Validated novel MAGs are then integrated into the classification framework. This involves: (1) converting GTDB taxonomy into the NCBI taxdump format, (2) assigning unique taxonomic identifiers for incomplete classifications, (3) rebuilding the Kraken2 database using kraken2-build with 32 threads. These augmented databases grow with each analysis and will be used as a long-term reference set for future projects.

### Phase 5: Functional Annotation

Functional annotation was accomplished using a multi-step approach. Prodigal (v2.6.3) [44] was employed in metagenomic mode (-p meta) for gene prediction from assembled contigs. For detailed functional assignments, eggNOG-mapper (v2.1.9) [45] was run in DIAMOND mode with in-memory database loading (--dbmem -- usemem) to compare predicted proteins to Clusters of Orthologous Groups (COG) [46] and Kyoto Encyclopedia of Genes and Genomes (KEGG) [47] orthology databases. The KEGG hierarchy database file (ko.txt) was used to set the levels of the KEGG pathways. The KEGG KO hierarchy file was retrieved using the KEGG REST API (http://rest.kegg.jp/get/br:ko00001). carbohydrate-active enzymes (CAZymes) annotation was performed using dbCAN3 (v4.1.0) [48] with HMMER as the core search algorithm (-t hmmer), supplemented by DIAMOND and eCAMI for exhaustive carbohydrate-active enzyme detection. Functional annotations were systematically extracted from eggNOG-mapper output files using targeted parsing.

### Phase 6: Phylogenetic Analysis

Phylogenetic analysis was done using a stepwise approach combining taxonomic classification, tree construction, and enhanced visualization. First, the taxonomic classification of MAGs was performed using GTDB-Tk (v2.1.1) in classify workflow mode against the Genome Taxonomy Database (GTDB) release 226 [49]. The next tree for the recovered MAGs was constructed using GTDB-Tk’s de_novo_wf workflow with the --skip_gtdb_refs parameter to exclude reference genomes and focus on study-specific MAGs. For tree visualization, R packages (ape [50], RColorBrewer [51]) and ggtree [52]/ggplot2 [53] libraries were used.

### Phase 7: Abundance Estimation

Kraken2 (v2.1.2) was used to classify the initial reads, and then Bracken [54] was used to correct read redistribution biases, generating abundance estimates at species (S) and genus (G) taxonomic levels using 150 bp read length parameters and a minimum threshold of 10 reads. Individual sample outputs were combined into unified abundance matrices using a custom integration script.

### Visualization and Statistical Analysis

Alpha diversity measures such as the Shannon diversity index, the Simpson diversity index, and observed taxonomic richness were computed for every sample. Bray-Curtis dissimilarity matrices with hierarchical clustering by the Ward linkage method were used in beta diversity analysis. Visualization of data was created with matplotlib v3.6.0 [55], seaborn v0.12.0 [56], and plotly v5.14.0 [57]. Network analysis used NetworkX v3.0 [58]. Output formats included quality control reports, assembly statistics, MAG quality plots (completeness vs. contamination), taxonomic composition charts, functional annotation heatmaps, and phylogenetic trees. All figures were saved at 300 dpi in publication-quality formats.

### Validation Dataset

Three metagenomic datasets were utilized to compare pipeline performance: human gut microbiomes (PRJNA553191, n=16) [17], lettuce rhizosphere (PRJNA763048, n=15) [18], and buffalo rumen (PRJNA718720, n=15) [19]. Raw sequencing data was downloaded via SRA Toolkit v3.0.3 [59]. Detailed metadata for these datasets is available in Tables S1, S2, and S3.

## Results

### MetaMAG Explorer Pipeline Implementation

We developed MetaMAG Explorer as a comprehensive computational pipeline for end-to-end metagenome-assembled genome (MAG) analysis with integrated database augmentation capabilities. From initial quality control to final abundance estimation, the pipeline architecture consists of seven sequential processing phases that can be operated separately or as an automated pipeline. These components are constructed as fifteen separate modular components (**Figure 1A**). MetaMAG Explorer’s unique feature is its novel MAG detection and database extension module **(Figure 1B)**, which automatically adds confirmed novel MAGs to customized Kraken2-compatible databases, verifies their novelty through dereplication against carefully curated collections, and systematically classifies unclassified genomes using GTDB-Tk classification. Three main reference collections (RUG, RGMGC, and MGnify) are used in the pipeline’s customized processing for rumen microbiome research. This approach establishes a feedback loop where every additional dataset enhances the reference database and improve future studies. The pipeline makes structured outputs that include high-quality MAGs with detailed quality reports, new genome sets with full taxonomic and functional annotations, better classification databases for better metagenomic profiling, and publication-quality visualizations for all the main analytical modules.

**Figure 1:**
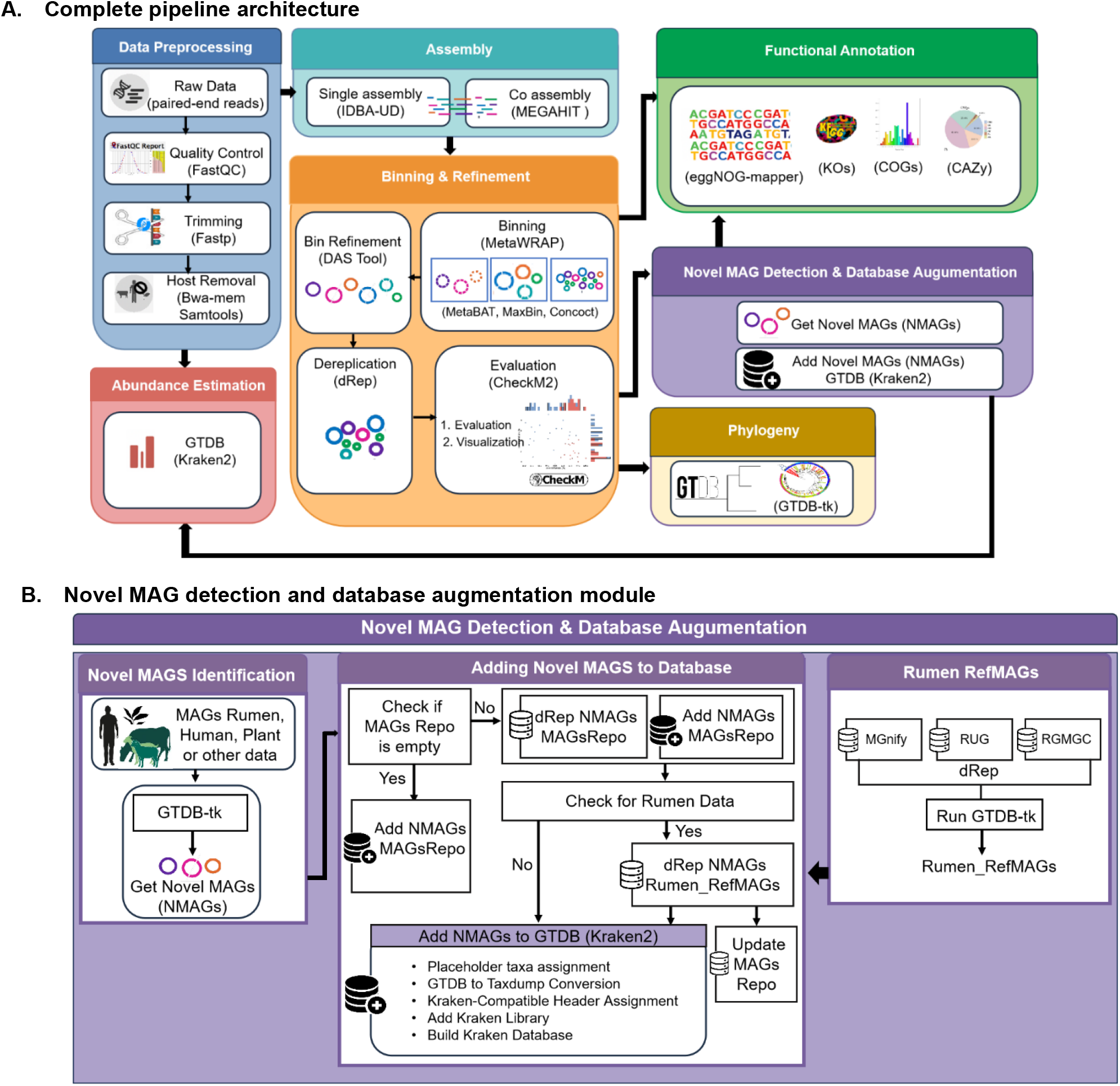
MetaMAG Explorer pipeline architecture and novel MAG detection workflow. (A) Pipeline overview. The workflow comprises modular stages including Data Preprocessing (quality control, trimming, and host read removal), Assembly (single-sample with IDBA-UD or co-assembly using MEGAHIT), Binning & Refinement (using MetaWRAP, DAS Tool, dRep, and CheckM2 for quality assessment and dereplication), Functional Annotation (using eggNOG-mapper to assign KOs, COGs, and CAZy families), Novel MAG Detection & Database Augmentation (identifying unclassified genomes and integrating them into custom Kraken2-compatible GTDB-based database), Phylogenetic Analysis via GTDB-Tk, and Abundance Estimation using Kraken2 classification. (B) Novel MAG detection and database augmentation workflow. The process consists of three key stages: identification of novel MAGs using GTDB-Tk classification; dereplication and integration into a shared MAG repository; and final formatting and augmentation of a custom Kraken2-compatible database, with a specialized process for MAGs of rumen origin that incorporates curated datasets (RUG, RMGMC, MGnify) for additional dereplication and validation.

### Performance Evaluation Across Diverse Metagenomic Datasets

To comprehensively examine the performance of MetaMAG Explorer, three test datasets from distinct environments are selected. These include (1) human gut microbiome samples (PRJNA553191, n=16), which were selected from the study that described microbial diversity across the human lifespan, including long-lived individuals such as centenarians and semi-supercentenarians [17]; (2) lettuce rhizospheric soil samples (PRJNA763048, n=15), from four cultivar types of cultivated lettuce (Lactuca sativa L.) and a bulk soil control in Gauteng, South Africa [18]; and (3) dairy Murrah buffalo rumen microbiota samples (PRJNA718720, n=15 samples); these samples were from four age groups [19]. This dataset was chosen to test how well the pipeline works across human, plant, and animal-associated microbiomes.

### Dataset Processing and Quality Control Performance

MetaMAG Explorer consistently performed well during preprocessing across all the samples. Post-quality control read retention was excellent: 93.9% for human gut microbiomes, 99.4% for lettuce rhizosphere soil, and 98.3% for rumen microbiomes. Host read removal was effective in all the datasets, with ∼26.9% of reads derived from humans being removed from gut samples, ∼4.1% of bovine sequences being removed from rumen samples, and ∼5.1% of plant-derived reads being removed from lettuce rhizosphere datasets. Assembly quality reflected the natural complexity of the community. Rhizosphere soil samples were more fragmented (median N50: 2,146 bp) but with higher taxonomic diversity, while human gut samples were more contiguous (median N50: 3,284 bp) and less fragmented, as would be anticipated given their lower taxonomic diversity. Rumen assemblies comprised intermediate contiguity (median N50: 2,792 bp) but also the largest aggregate assembly sizes (e.g., 295 ± 21 Mb), reflecting the metabolic and genomic richness of the rumen microbiota.

### Comprehensive MAG Recovery and Quality Assessment

From all three datasets, a total of 233 MAGs were recovered, of which 121 achieved high-quality (≥90% completeness, ≤5% contamination) status as determined by the CheckM2 evaluation. MAGs from the human gut showed the highest completeness (93.1 ± 5.1%) and the lowest contamination (3.1 ± 2.8%), reflecting the rich and strain-diverse nature of the human gut microbiome [2, 3]. MAGs from the rhizosphere displayed similar completeness (93.1 ± 5.8%) but showed slightly increased contamination levels (4.1 ± 2.8%), indicating the richness and variety of the soil microbial community. Rumen MAGs exhibited a somewhat lower average completeness (89.8 ± 5.3%) and moderate contamination (3.5 ± 2.4%), reflecting their genetic richness and complexity (**Figures 2A and 2B**). **Supplementary Figures S2, S3, and S4** provide details of genome completeness and contamination. Taxonomic profiling of 233 classified MAGs revealed environment-specific patterns of microbial diversity across 12 bacterial and 1 archaeal phylum. Human gut MAGs were enriched and dominated by *Bacillota* (66% of classified MAGs) and *Actinomycetota* (19%), consistent with the *Firmicute*-dominated composition characteristic of human gut microbiomes [2]. MAGs from plant rhizosphere were dominated by *Pseudomonadota* (60% of plant MAGs). Rumen MAGs, on the other hand, have balanced representation of *Bacteroidota* (36%) and *Bacillota* (30%), and specialist lineages like methanogenic Archaea (*Methanobacteriota*) and fiber-degrading *Spirochaetota*. This diversity shows the rumen’s advanced metabolic potential (**Figures 2C and 2D**).

**Figure 2:**
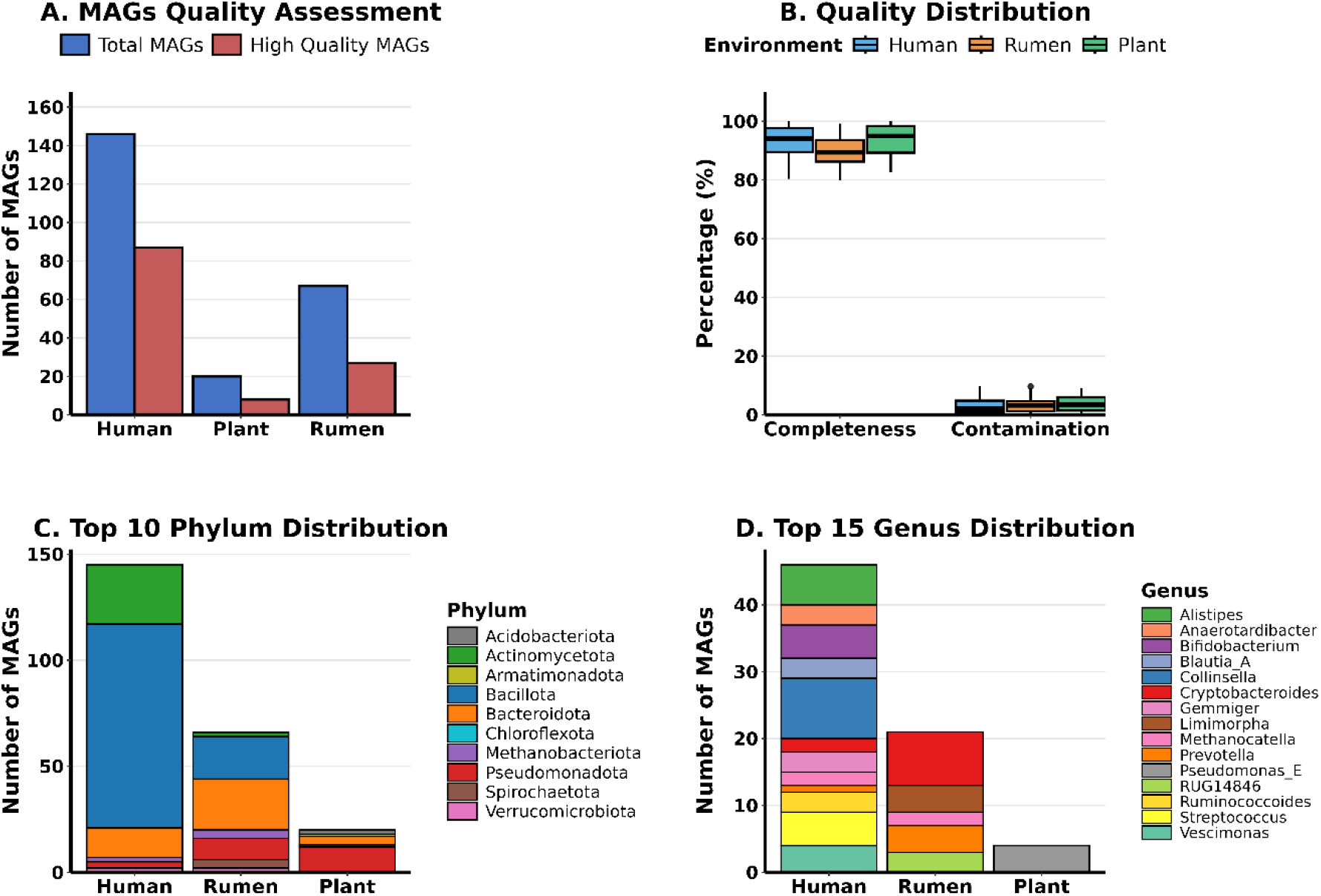
MAG quality and taxonomic diversity across microbiome environments. **(A)** Total and high-quality MAG counts per environment. **(B)** Genome quality distributions showing completeness and contamination percentages. **(C)** Top 10 phyla and **(D)** top 15 genera based on MAG abundance. High-quality MAGs were defined as ≥90% complete with ≤5% contamination. Numbers represent MAG counts: Human (n=146), Rumen (n=67), Plant (n=20).

### Novel MAG Discovery and Taxonomic Characterization

48 novel MAGs were recovered across all datasets, constituting 20.0% of the total high-quality MAGs (**Figure 3A**). These genomes were categorized as novel based on stringent criteria, including dereplication, taxonomic uniqueness, and validation against public reference databases. The quality of these genomes is high, with over 80% completeness and less than 10% contamination (**Figure 3B**). Distribution of novel MAGs was strongly environment dependent. The highest number of these MAGs are from rumen samples (58.3%, n = 28) with species from *Bacteroidota, Spirochaetota, Bacillota*, and *Methanobacteriota*. Plant rhizosphere samples contribute to the second most novel MAGs (29.2%, n = 14), including predominant representation from *Pseudomonadota* and *Acidobacteriota*. Whereas from human gut samples, although the highest number of MAGs were recovered (146 MAGs), they only contributed to 6 novel MAGs (12.5%), with most of these being from *Actinomycetota*, a genus underrepresented in existing human gut reference genomes. This distribution aligns with the known bias toward human-associated microbiomes in genomic databases [2, 3] compared to environmental samples [8, 37]: rumen and soil environments are still under sampled, whereas human gut microbiomes have been thoroughly studied. Taxonomic profiling illustrates discrete community structure by environment. Genus-level taxonomic assignments identified environment-specific patterns, such as enrichment of *RUG1846, Colidextribacter*, and *Cryptobacteroides* within rumen MAGs, while plant-derived MAGs harbored taxa such as *Caulobacter* and other poorly characterized genera. Human-associated MAGs, although few, included phylogenetically distinct taxa that may represent novel clades of formerly well-characterized groups. Overall, these findings emphasize the environmental uniqueness and genomic novelty that are made available through MetaMAG Explorer’s recovery pipeline (**Figures 3C and 3D**).

**Figure 3:**
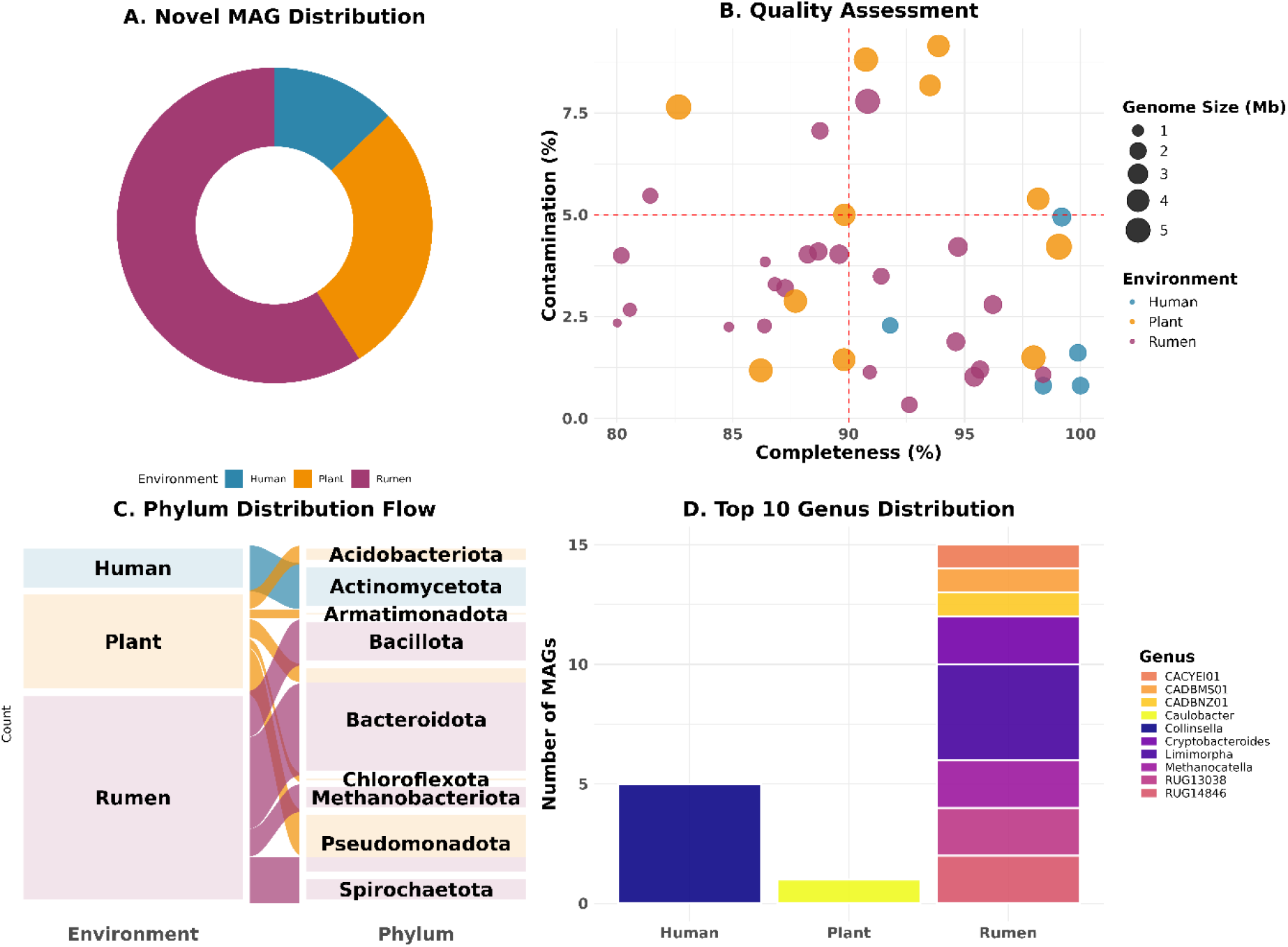
Novel metagenome-assembled genome discovery and taxonomic characterization across microbiome environments. (A) Distribution of 39 novel MAGs across human gut, rumen, and plant rhizosphere microbiomes, with rumen samples contributing to the majority, followed by plant rhizosphere and human gut. (B) Quality assessment showing genome completeness versus contamination levels, with circle sizes representing genome size (Mb). Dashed red lines indicate high-quality thresholds (≥90% completeness, ≤5% contamination). (C) Alluvial flow diagram illustrating phylum-level taxonomic distribution, with flow thickness representing the number of MAGs transitioning from each environment to phylogenetic classification. (D) Distribution of the top 10 most abundant genera among novel MAGs across the three environments.

### Comprehensive Functional Characterization of Novel MAGs

MetaMAG Explorer conducted automated functional characterization of all 48 novel MAGs utilizing the dbCAN, KEGG, COG, and eggNOG-mapper databases. Here we are showing the results from rumen MAGs due to the highest recovery rate (28 of 48). Results for human gut (n=6) and plant rhizosphere (n=14) are included in Supplementary Materials (**Figures S5 and S6**). The CAZyme annotations showed that glycosyltransferases (GT) and glycoside hydrolases (GH) were the most abundant, which supports the idea of the rumen’s role in the degradation of complex plant carbohydrates. Among these, *Bacteroidota* MAGs (n=12) had the highest GH and CE abundance, while *Bacillota* (n=7) showed varied enzyme patterns. Carbohydrate-binding modules (CBM) were found in all MAGs, showing their ability to bind to complex substrates (**Figure 6A**). The COG functional analysis indicated that genes are spread across different taxa (**Figure 6B**). The KEGG pathway flow represents metabolism as the main category (**Figure 6C**). Further, the detailed flow for the metabolism category was observed.

**Figure 6:**
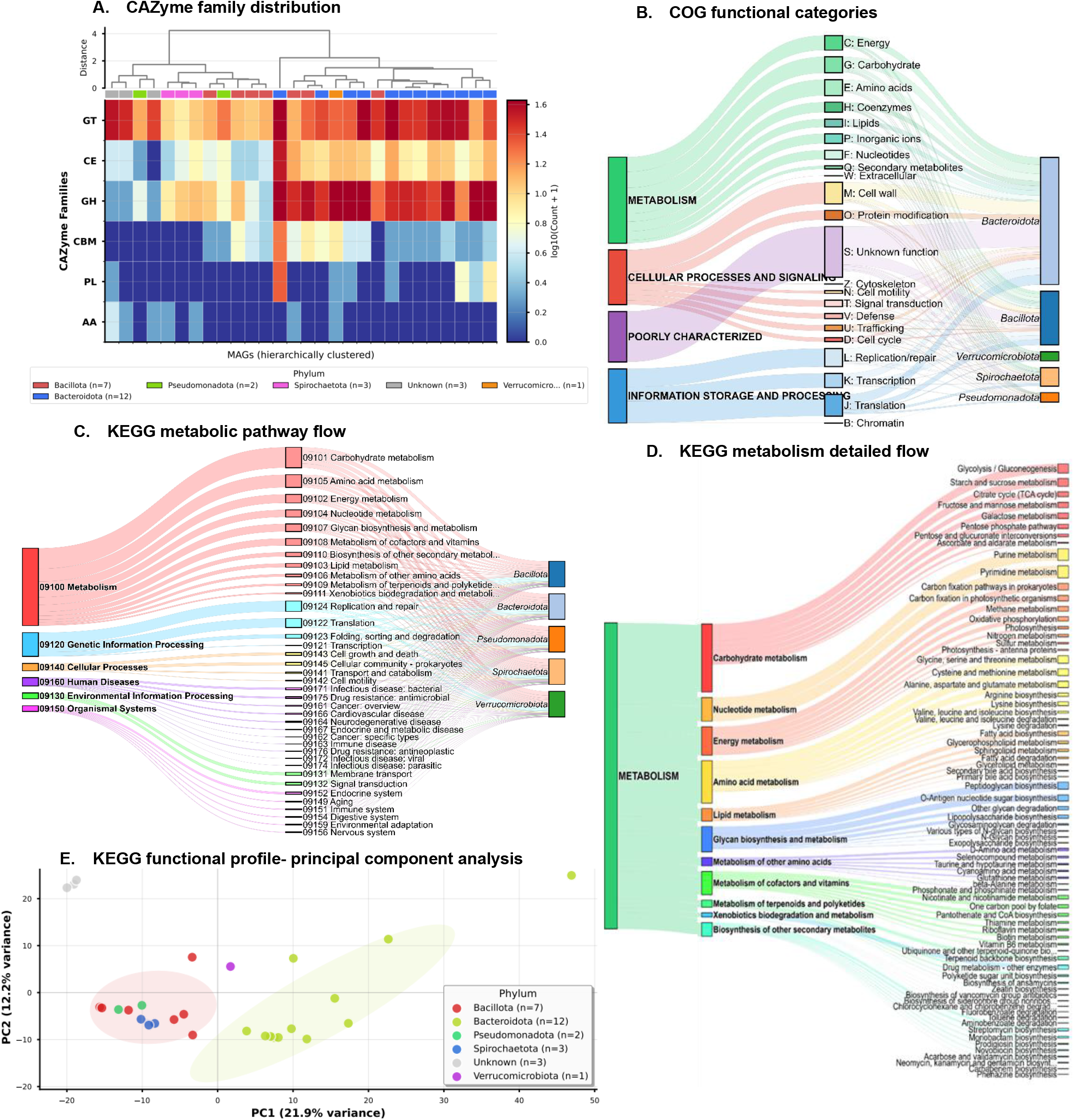
Functional characterization of novel MAGs from rumen samples. **(A)** Distribution of carbohydrate-active enzyme (CAZyme) families across novel MAGs, clustered by genome and colored by phylum, with major classes including glycosyltransferases (GT), carbohydrate esterases (CE), glycoside hydrolases (GH), carbohydrate-binding modules (CBM), polysaccharide lyases (PL), and auxiliary activities (AA). **(B)** COG functional category assignments for predicted proteins, with the percentage of functions related to metabolism, cellular processes, information storage, and poorly characterized proteins. **(C)** KEGG metabolic pathway flow, connecting high-level functional categories such as metabolism, cellular processes and others to the taxonomic groups in which they are found. **(D)** Breakdown of individual pathways within the “Metabolism” category. **(E)** Principal component analysis (PCA) of KEGG functional profiles, with clustering of MAGs by phylum and good separation of functional traits between taxonomic groups.

The Sankey diagram illustrates metabolism (green) as the predominant category, with flows directed towards glucose metabolism, amino acid metabolism, and energy production. The next dominant group was cellular processes, followed by information storage. Unknown functions were rare. *Bacteroidota* and *Bacillota* were involved in more functions, while the *Pseudomonadota, Spirochaetota*, and *Verrucomicrobiota* had more specialized profiles. KEGG pathway mapping revealed the broad metabolic potential for these microbes (**Figure 6D**). The citrate cycle, glycolysis/gluconeogenesis, and other pathways of sugar metabolism feed into the prevailing category of carbohydrate metabolism (red) based on the Sankey diagram. Lipid metabolism and glycan biosynthesis showed smaller inputs, while energy metabolism, nucleotide metabolism, and amino acid metabolism were all well-represented (yellow, orange, and yellow). This metabolic breadth indicates the functional completeness of rumen MAGs. Principal component analysis revealed a phylogenetic grouping of functional traits (**Figure 6E**). There was a total of 34.1% variance explained by PC1 (21.9%) and PC2 (12.2%). The *Bacteroidota* MAGs (yellow, n=12) formed their own cluster on the right, while the *Bacillota* MAGs (red, n=7) clustered on the left. The fact that the remaining phyla, such as *Bacillota* MAGs (red, n=7), *Pseudomonadota* (n=2), and *Spirochaetota* (n=3), have intermediate places indicates that rumen MAG functional potential is substantially influenced by phylogeny.

The functional links between MAGs were found by network analysis (**Figures S7 and S8**). The members of the *Bacteroidota* (yellow) and *Bacillota* (red) MAGs showed phylogenetic clustering in the KEGG network, where they formed separate groups connected by shared pathways (K07133, K03088, and K02004 being the most common). All MAGs demonstrated markedly denser connectivity with extensive edges in the CAZyme network, especially those that included the two primary enzyme categories, glycoside hydrolases (GH) and glycosyltransferases (GT). The sparse KEGG network shows phylogenetically constrained metabolic pathways, while the CAZYme-dense network suggests possible synergistic interactions in plant biomass degradation. The pipeline showed how MetaMAG Explorer can perform thorough functional analysis across various microbiome environments by producing all visualizations automatically.

### Phylogenetic Placement of Recovered MAGs

MetaMAG Explorer builds phylogenetic trees for all recovered MAGs using GTDB-Tk. By placing our MAGs onto the GTDB-Tk reference tree, we could see where they belong in the bacterial family tree and identify novel species. These new MAGs were spread across many different phyla, showing that new diversity is not limited to one group but is widely distributed in the microbiome. The trees, shown in both rectangular and circular views, give a clear picture of the taxonomic makeup of our genomes (**Figure 7A, 7B**). Many novel MAGs appear next to well-known lineages, suggesting they are new branches of existing families. When we added functional data to the circular tree, we found that novel MAGs often share the same key traits as their closest relatives; for example, fiber-degrading groups are rich in glycoside hydrolases (**Figure 7C**). Overall, the phylogenetic analysis showed that the new MAGs are taxonomically diverse and functionally significant. This wide distribution and clear functional roles suggest that these new genomes are important members of their communities and will improve how we classify and understand microbiomes. Comparable analyses for the plant-associated and human-associated datasets are provided in (**Figures S9 and S10**), respectively.

**Figure 7:**
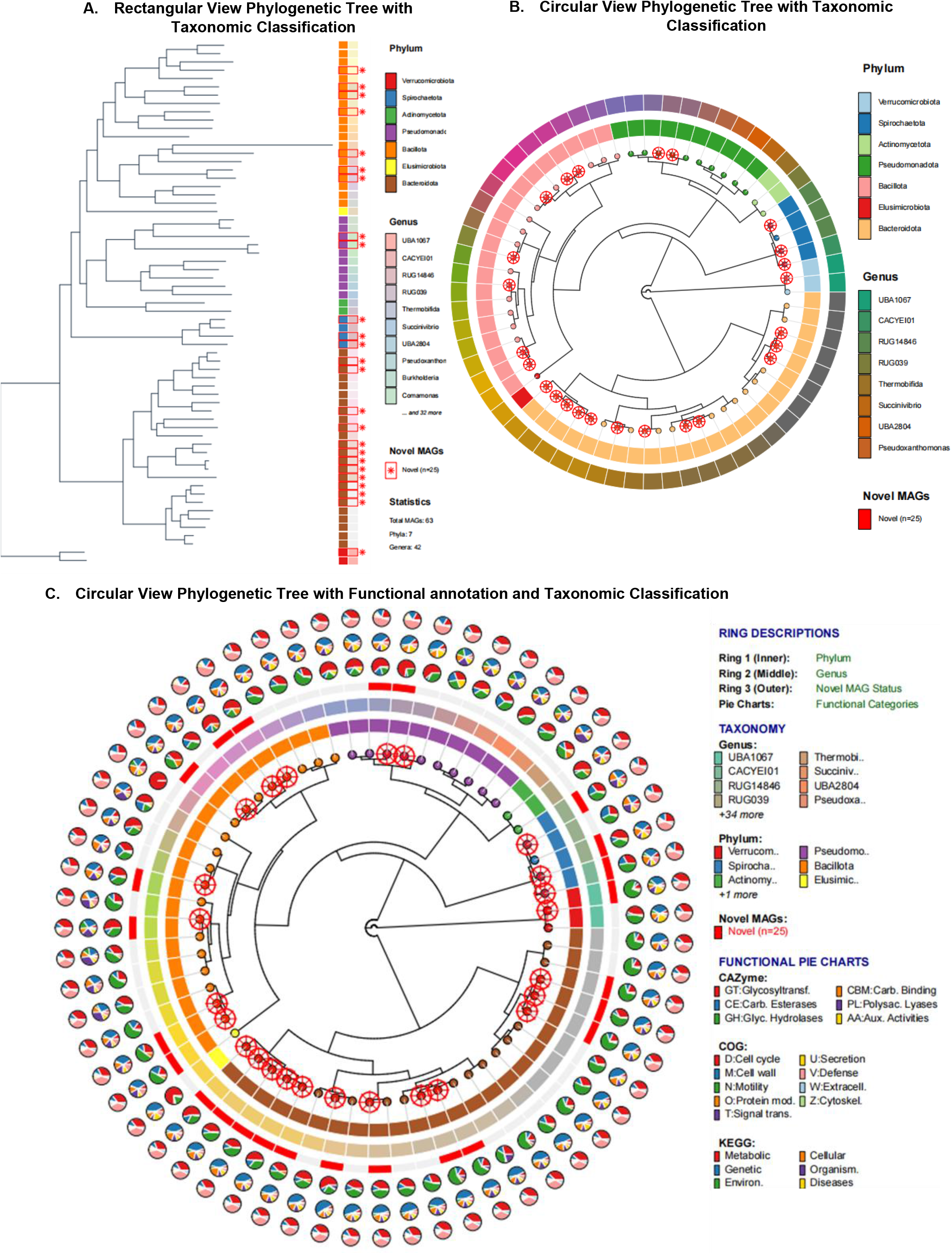
Phylogenetic placement and taxonomic classification of novel MAGs. **(A)** A rectangular phylogenetic tree with taxonomic classification at the phylum and genus levels; new MAGs are marked in red. **(B)** A circular phylogenetic tree that shows taxonomic classification by phylum (the outer ring) and genus (the inner ring), with new MAGs marked in red. **(C)** A circular phylogenetic tree that combines functional annotation with taxonomic classification. The circular phylogenetic tree shows different annotation layers, from the inner ring to the outer ring. The phylum classification for each MAG is shown in Ring 1 (the innermost ring). The second ring (middle) shows the genus classification. Ring 3 (the outer solid bars) shows that the MAG is new, and the red bars show genomes that have been found to be new. There are three concentric functional annotation layers outside of these rings. The innermost functional ring shows CAZyme classes, the middle functional ring shows COG functional categories, and the outermost functional ring shows KEGG pathway categories. The pie charts around these functional categories show how they relate to each MAG.

### Abundance estimation with updated Kraken2 database

To demonstrate the impact of database augmentation, we focused on plant rhizosphere samples here. Soil and rhizosphere environments are severely under sampled in current databases [4, 8], so this is where we expect to find the value of the novel MAGs. When we included the new MAGs in the Kraken2 database, several were among the top 20 most abundant species, namely positions 6, 8, and 12, each comprising 0.5-0.8% of the community (**Figure 8A**). Their presence in all plant samples showed that they are prominent members of the rhizosphere microbiome, not rare species. Soil microbiomes have been studied for decades, yet organisms comprising nearly 1% of the community have remained undetected. These are likely to play important ecological roles that were completely missed before. Previous studies of plant rhizosphere has just labeled these as unclassified. The Shannon index and PCA, which are measures of diversity, show that including them gives a clearer and more accurate picture of community composition (**Figure 8B**). The classification improvement for these samples was very minute, but it shows that major species that were completely hidden in previous analyses have been identified here. This shows how MetaMAG Explorer can find important microbes in environments that haven’t been studied as much.

**Figure 8:**
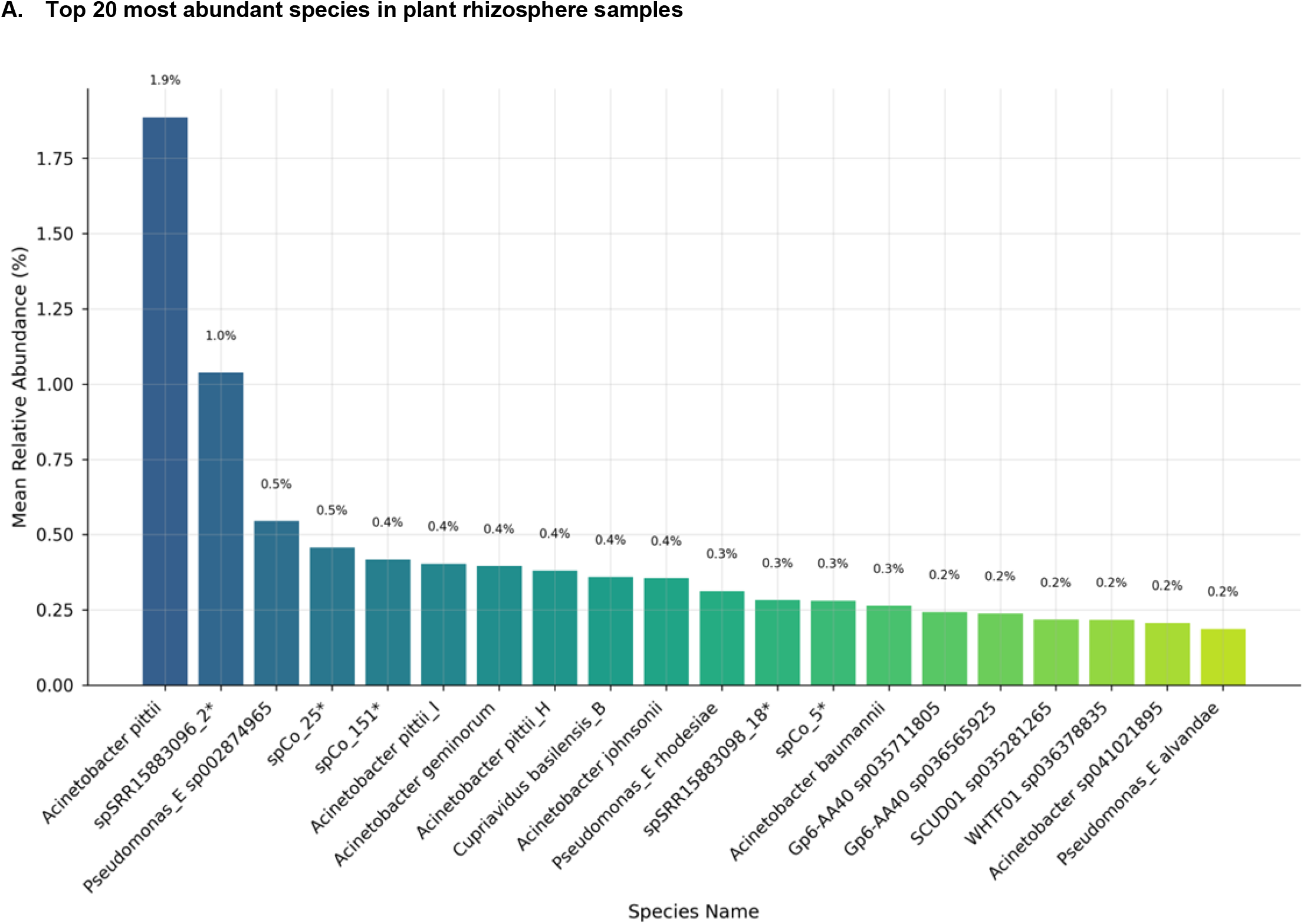
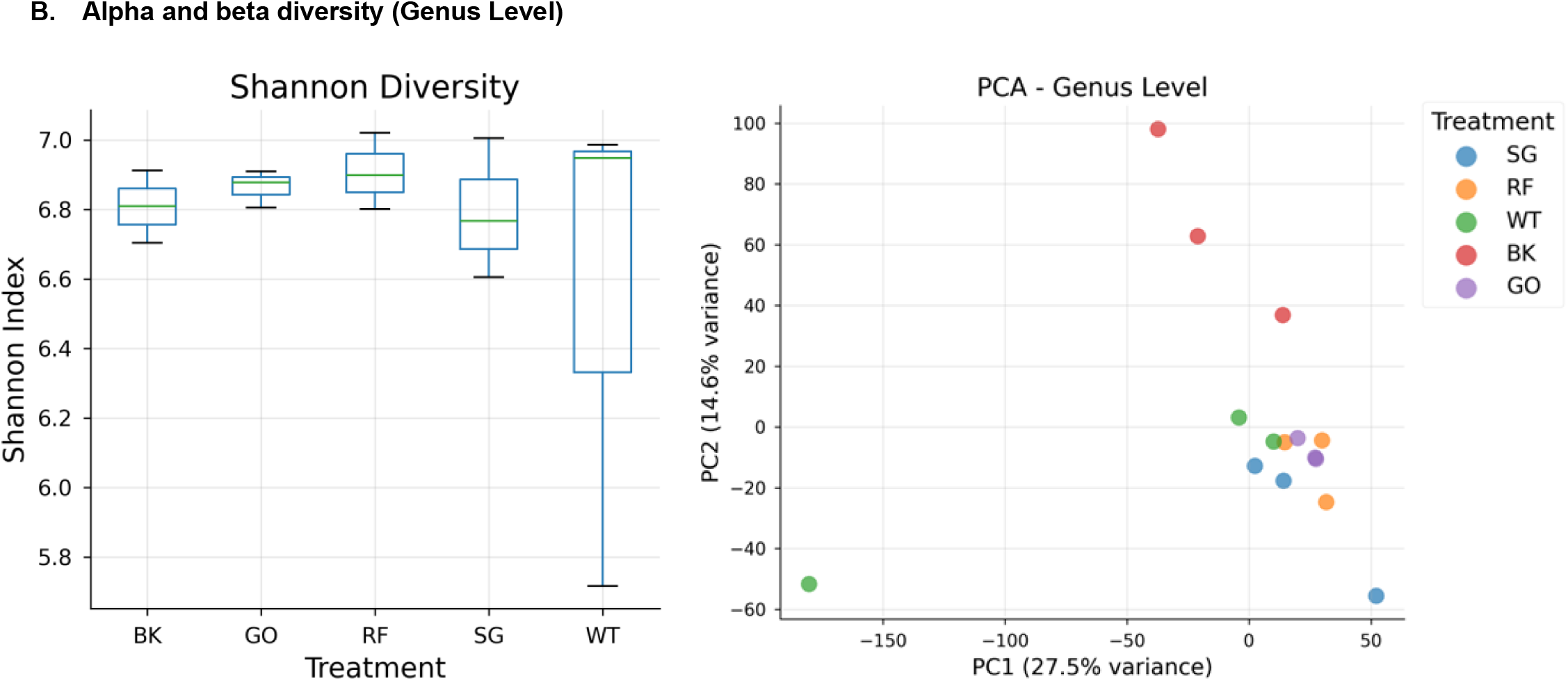
Diversity and composition of microbial communities in plant rhizosphere samples. The top 20 most prevalent species are displayed in a bar plot along with the mean relative abundance (%) for each sample. With relative abundance percentages on the y-axis ranging from 0% to approximately 1.8%. (B) Diversity in alpha and beta at the genus level. When comparing diversity across treatments (BK, GO, RF, SG, and WT), the Shannon diversity index (left) shows that diversity is generally high with only minor variation between treatments. Based on genus-level community composition, Principal Component Analysis (PCA, right) reveals clustering patterns that represent variations in the structure of the microbial communities across treatments.

### Impact of Database Augmentation on Classification

While the rhizosphere results showed how even a few novel MAGs can identify important microbial communities in under-sampled environments, the rumen dataset lets us see the impact when many new genomes are recovered. The rumen data shows another important point: adding new genomes does not just slightly increase the percentage of classified reads, it also improves the accuracy of where those reads end up. Adding these 28 novel MAGs from this study and 908 additional rumen reference MAGs to the Kraken2 database allowed us to classify 5.4 million additional reads across 15 samples, raising classification rates by an average of 1.27% (**Table 1**). A 1-2% gain might seem small at first, but here is the key: First, these 28 new MAGs were identified in every sample, indicating a 100% prevalence. These are not rare organisms that could be technical noise; they are stable community members that are found in all animals. Second, 1.27% (5.4 million) reads now have taxonomic classification. Third, consider the compounding effect by accumulating novel MAGs from ongoing projects and diverse sources. Further, if we see the reads assigned to novel MAGs, it is apparent that database augmentation reassigns reads that were previously classified at higher or incorrect levels to the more specific and ecologically relevant novel genomes in addition to classifying previously unclassified reads. This means that the right novel MAGs are now assigned to hundreds of thousands of reads per sample that were previously categorized at the genus, family, or even unrelated species level. This reallocation is evident from the global breakdown (**Figure S11**), where an average of approximately 2.0 million reads per sample were redistributed to the augmented genome set, with approximately 17.3% going to the novel MAGs and 82.7% going to the additional rumen reference MAGs.

**Table 1:**
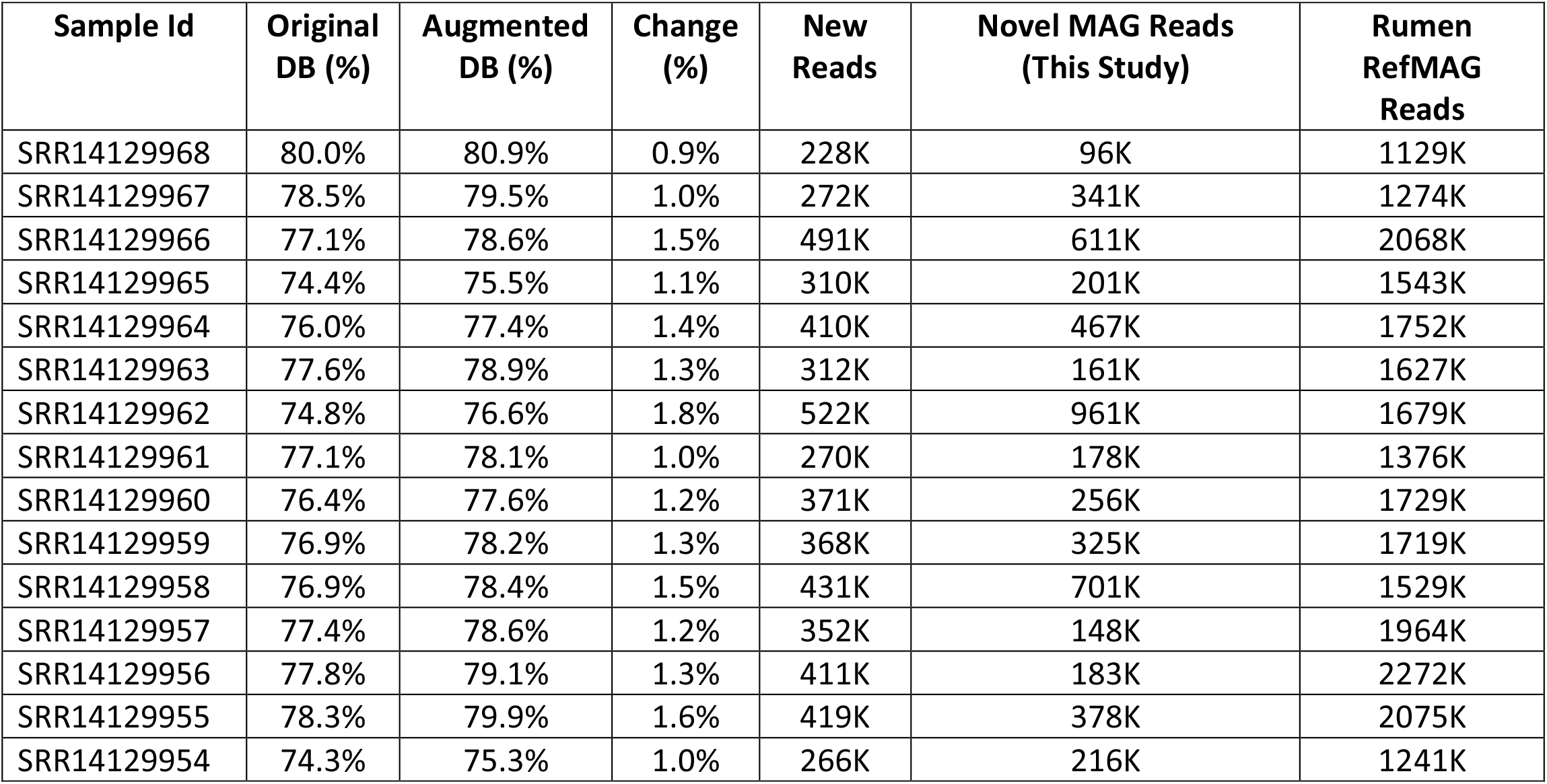
Database augmentation’s effect on rumen metagenome read classification. The original and enhanced Kraken2 databases’ classification performance for 14 rumen samples is contrasted in the table. The percentage of reads classified using the original database (Original DB %), the percentage classified following augmentation with novel and reference rumen MAGs (Augmented DB %), and the net change in classification percentage (Change %) are displayed in the columns Additionally displayed for each sample is the quantity of newly classified reads (New Reads). The table shows the number of reads specifically attributed to the 28 novel MAGs recovered in this study (Novel MAG Reads) and to the 908 additional rumen reference MAGs integrated from public catalogs (Rumen RefMAG Reads) to differentiate the contribution of various genome sets. Augmentation increased classification rates in all samples, and both reference and novel MAGs significantly increased read reassignment.

## Discussion

MetaMAG Explorer demonstrates that dynamic database augmentation through automated integration of project-specific MAGs can substantially improve metagenomic classification accuracy. The pipeline’s iterative approach, recovering MAGs, validating novelty, and integrating them into Kraken2-compatible databases, improved classification and correctly assigned previously unclassified reads.

MetaMAG Explorer addresses critical gaps that existing pipelines have not resolved. Other pipelines like SqueezeMeta, ATLAS, and METABOLIC provide complete MAG recovery and analysis utilities, but they treat recovered MAGs as endpoints rather than resources for database enhancement. EasyMetagenome offers user-friendly, rich-pipeline visualization but similarly lacks database augmentation capabilities. In contrast, MetaMAG Explorer provides the feature of automatic identification, validation, and integration of novel MAGs into Kraken2-compatible databases, continuously improving classification accuracy. Additionally, MetaMAG Explorer includes specialized modules for rumen microbiome analysis with curated reference datasets (RUG, RGMGC, MGnify), addressing the unique challenges of this under-sampled environment.

In the example dataset we found 48 novel high-quality MAGs across three test environments with 100% prevalence in their respective environments. This highlights not only the extensive microbial diversity absent from current references but also that these gaps represent stable, functionally vital organisms rather than rare or marginal species. The distribution reflects database biases: human gut samples contributed to most of the overall MAGs (146); they only contributed 12.5% of novel genomes, while rumen samples contributed 58.3% of novel genomes despite making up only 29% of total MAGs. This discrepancy is consistent with the thorough description of gut microbiomes in Western humans rather than environmental systems. Adding these new genomes to databases that work with Kraken2 had immediate benefits. For example, it improved classification rates, as shown by the large number of read reassignments in rumen datasets and the discovery of ecologically important, previously unclassified groups in rhizosphere communities.

Functional analysis offered additional perspectives on the value of the new genomes. In the rumen, members of *Bacteroidota* and *Bacillota* carried many CAZymes, enzymes that specialize in breaking down complex carbohydrates. These groups are well suited to this environment, where efficient fiber degradation is essential. Their recovery shows how MetaMAG Explorer can reveal functional groups that play central roles in ecosystem processes. Although the overall increase in classification rate was modest, many reads were reassigned from misclassified groups to these novel MAGs. Over time, this iterative approach strengthens the reference library, making it more reliable for long-term microbiome studies.

This approach for self-improvement has a big effect on how metagenomic studies are carried out. In MetaMAG Explorer, projects are not treated as isolated studies. Instead, it supports a community-based approach, where each study adds to a growing body of knowledge. This is specifically important in under-sampled environments where such database integration helps to draw meaningful information. The cumulative impact of iterative improvement, whereby each study increases classification accuracy for subsequent jobs, can effectively decrease the percentage of unclassified reads over time. In applied contexts, such as enhancing the efficiency of the rumen microbiome to mitigate methane emissions or identifying beneficial taxa for agriculture, such advances in categorization have the potential to reveal novel biological targets or indicators that were previously overlooked.

## Conclusions

MetaMAG Explorer is a scalable pipeline that advances metagenomic research by automatically adding newly recovered MAGs to classification databases, allowing taxonomic resolution to improve with each analysis. By revealing previously unrepresented but ecologically significant genomes from diverse environments, it helps close gaps in existing reference resources. This self-updating approach is especially useful in environments that are still poorly studied, where incomplete databases have made biological interpretation difficult. In the future, MetaMAG Explorer will provide scientists with an expanding and self-improving framework, making it easier to connect novel genomes with their ecological roles and functional significance.

## Data Availability

All sequencing datasets analyzed in this study are publicly available from the NCBI Sequence Read Archive (SRA). Human gut microbiome data are available under accession PRJNA553191, lettuce rhizosphere under PRJNA763048, and buffalo rumen under PRJNA718720.

The MetaMAG Explorer pipeline is freely available at [https://github.com/msatti123/MetaMAG_Explorer]. We have developed comprehensive documentation including detailed HTML web pages that provide step-by-step installation instructions, user guides, parameter descriptions, and troubleshooting tips, all hosted as GitHub Pages at [https://msatti123.github.io/MetaMAG_Explorer/. The repository includes all source code, example datasets, test scripts, and configuration templates.

## Ethics Statement

This study analyzed previously published and publicly available sequencing datasets. No new human or animal samples were collected. Ethical approval and informed consent were obtained in the original studies [PRJNA553191, PRJNA763048, PRJNA718720]. Therefore, no additional ethical approval was required for the analysis presented here.

## Acknowledgements

We are grateful to the funding bodies, Milk Levy Fund (Mælkeafgiftsfonden), Aarhus, Denmark and Innovation Fund Denmark, which supported our research.

## Author Contributions

Maria Altaf Satti developed the MetaMAG Explorer pipeline, performed the analyses, interpreted the results, and wrote the initial manuscript draft. Zexi Cai conceived and designed the study, supervised the study, provided critical guidance, secured the funding, provided necessary resources, and contributed to reviewing and revising the manuscript. Both authors read and approved the final manuscript.

