## Supplementary_Figures for "MetaMAG Explorer: A Database-Augmenting Pipeline for Genome-Resolved Metagenomics and Enhanced Microbial Classification"

#### Figure S1

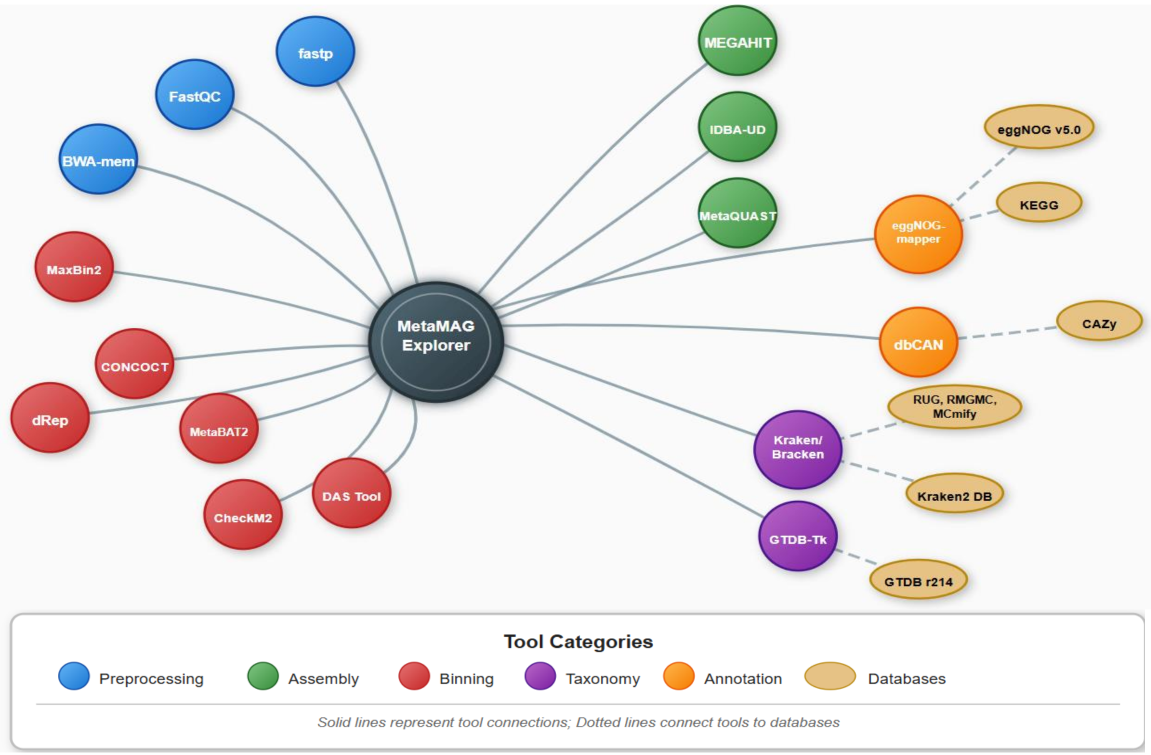


**Figure S1**: Tools and reference databases used in the MetaMAG Explorer pipeline. Nodes are color-coded by function: preprocessing (blue), assembly (green), binning and refinement (red), annotation (orange), and classification/phylogeny (purple). Dashed lines indicate associated reference databases.

#### Figure S2

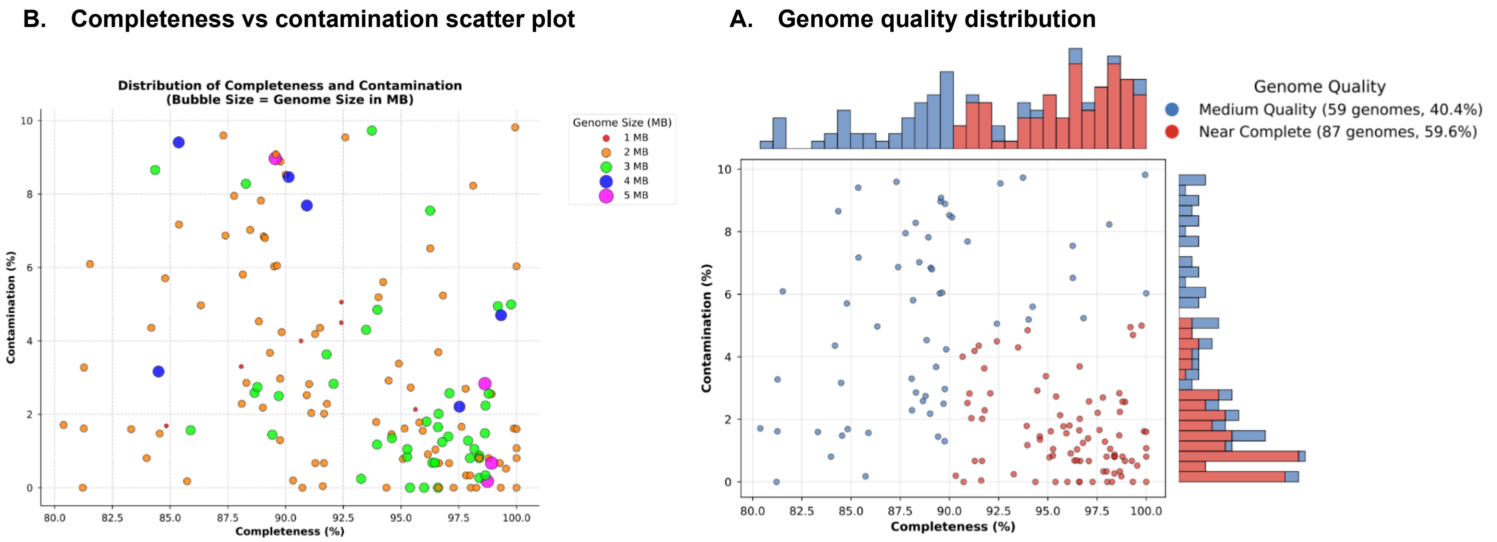


**Figure S2:** Quality assessment based on completeness and contamination for MAGs recovered from human samples. **(A)** Distribution of genomes by completeness, contamination, and genome size from 146 recovered MAGs, showing high completeness and low contamination that demonstrates the high caliber of genome reconstruction. **(B)** Quality group classification with accompanying histograms, revealing that 59 MAGs (40.4%) are of medium quality and 87 MAGs (59.6%) are near-complete.

### Figure S3

**A. Completeness vs contamination scatter plot**

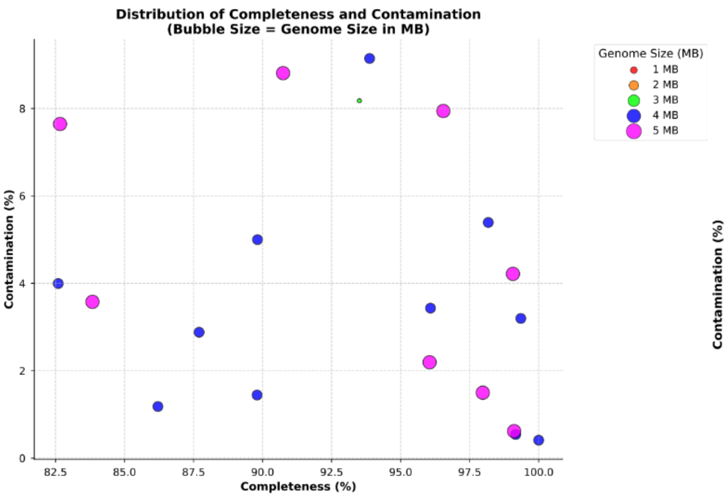


**B. Genome quality distribution**

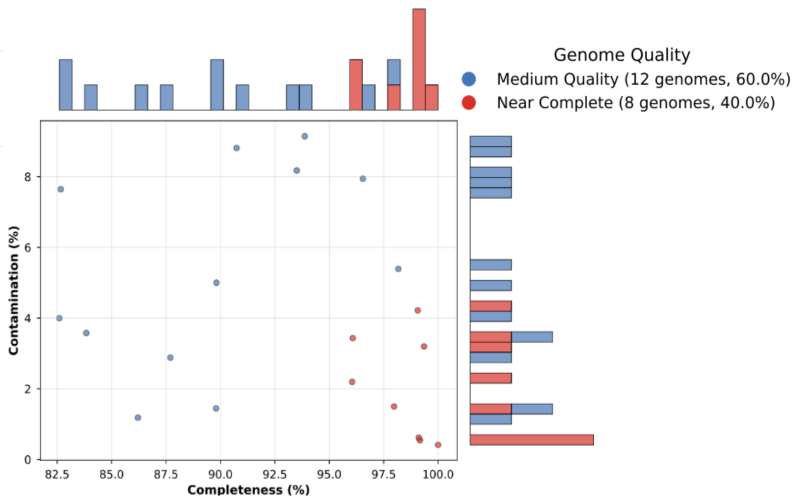


**Figure S3:** Quality assessment based on completeness and contamination for MAGs recovered from plant samples. (A) Distribution of genomes by completeness, contamination, and genome size from 20 recovered MAGs, showing high completeness with low contamination that indicates good genome reconstruction. (B) Quality group categorization with accompanying histograms, revealing that 12 MAGs (60.0%) are of medium quality and 8 MAGs (40.0%) are near-complete.

### Figure S4

**A. Completeness vs contamination scatter plot**

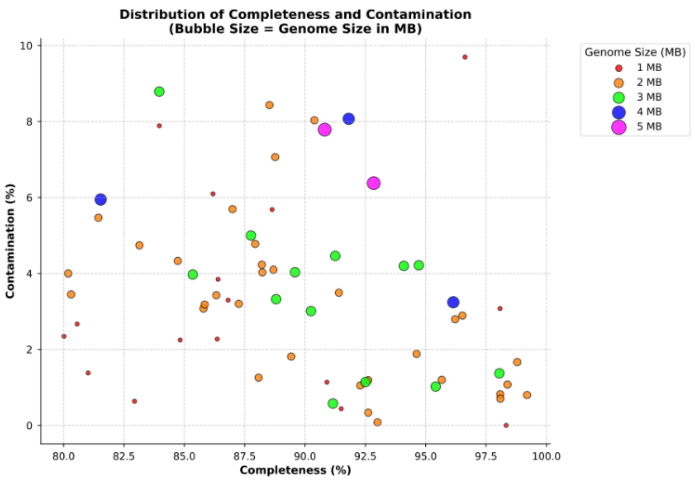


**B. Genome quality distribution**

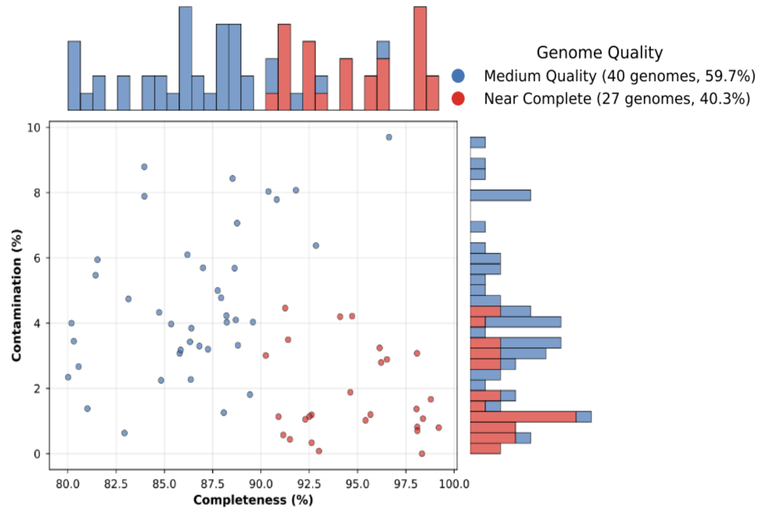


**Figure S4:** Quality assessment based on completeness and contamination for MAGs recovered from rumen samples. (A) Distribution of genomes by completeness, contamination, and genome size from 67 recovered MAGs, showing high completeness with low contamination that reflects reliable genome reconstruction. (B) Quality groupings with accompanying histograms, revealing that 40 MAGs (59.7%) are of medium quality and 27 MAGs (40.3%) are near-complete.

#### Figure S5

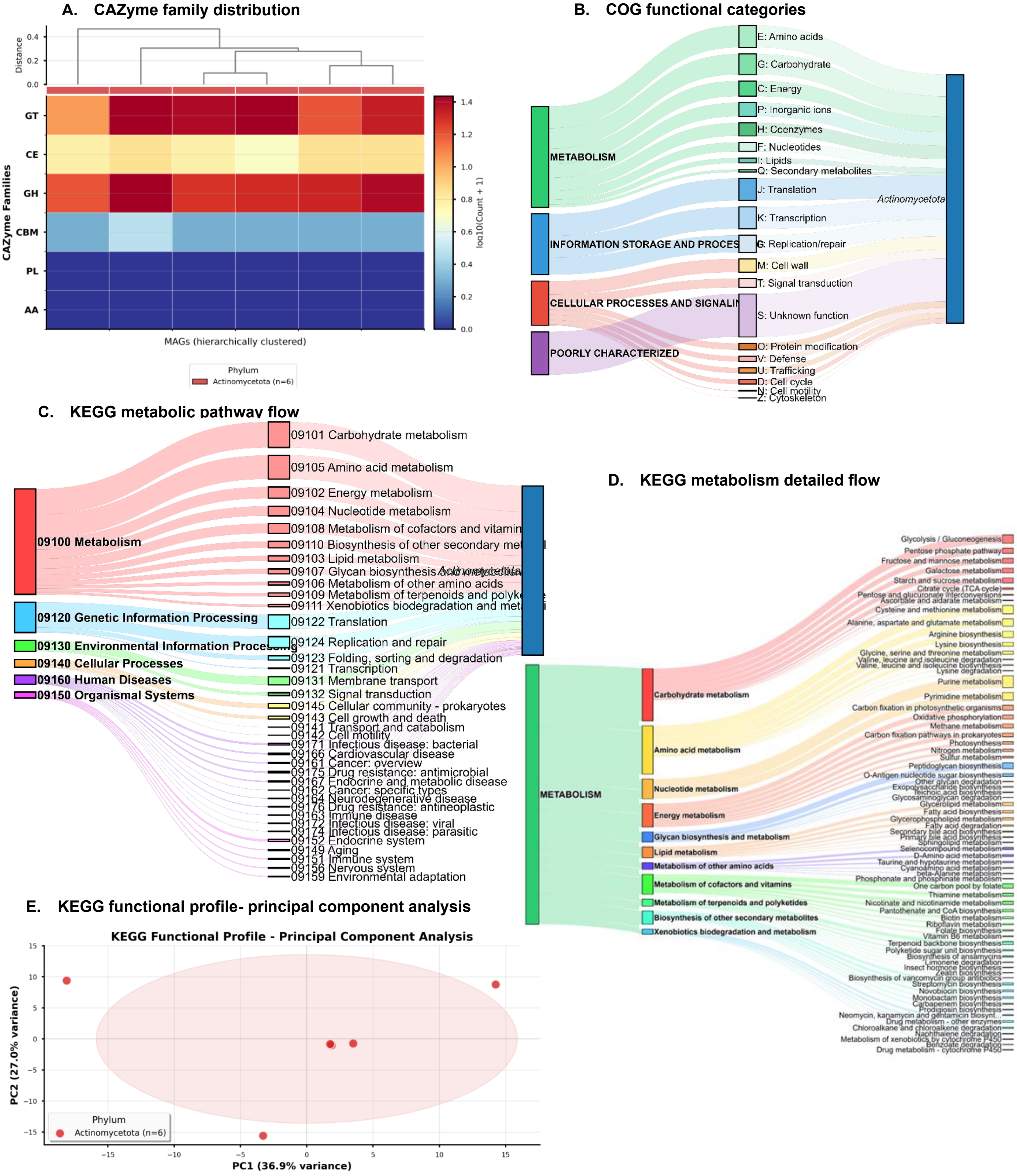


**Figure S5:** Functional characterization of novel MAGs from human. **(A)** Distribution of carbohydrate-active enzyme (CAZyme) families across novel MAGs, clustered by genome and colored by phylum. **(B)** COG functional category assignments for predicted proteins, with the percentage of functions related to metabolism, cellular processes, information storage, and poorly characterized proteins. **(C)** KEGG metabolic pathway flow, connecting high-level functional categories such as metabolism, cellular processes and others to the taxonomic groups in which they are found. **(D)** Breakdown of individual pathways within the "Metabolism" category. **(E)** Principal component analysis (PCA) of KEGG functional profiles, with clustering of MAGs by phylum.

### Figure S6

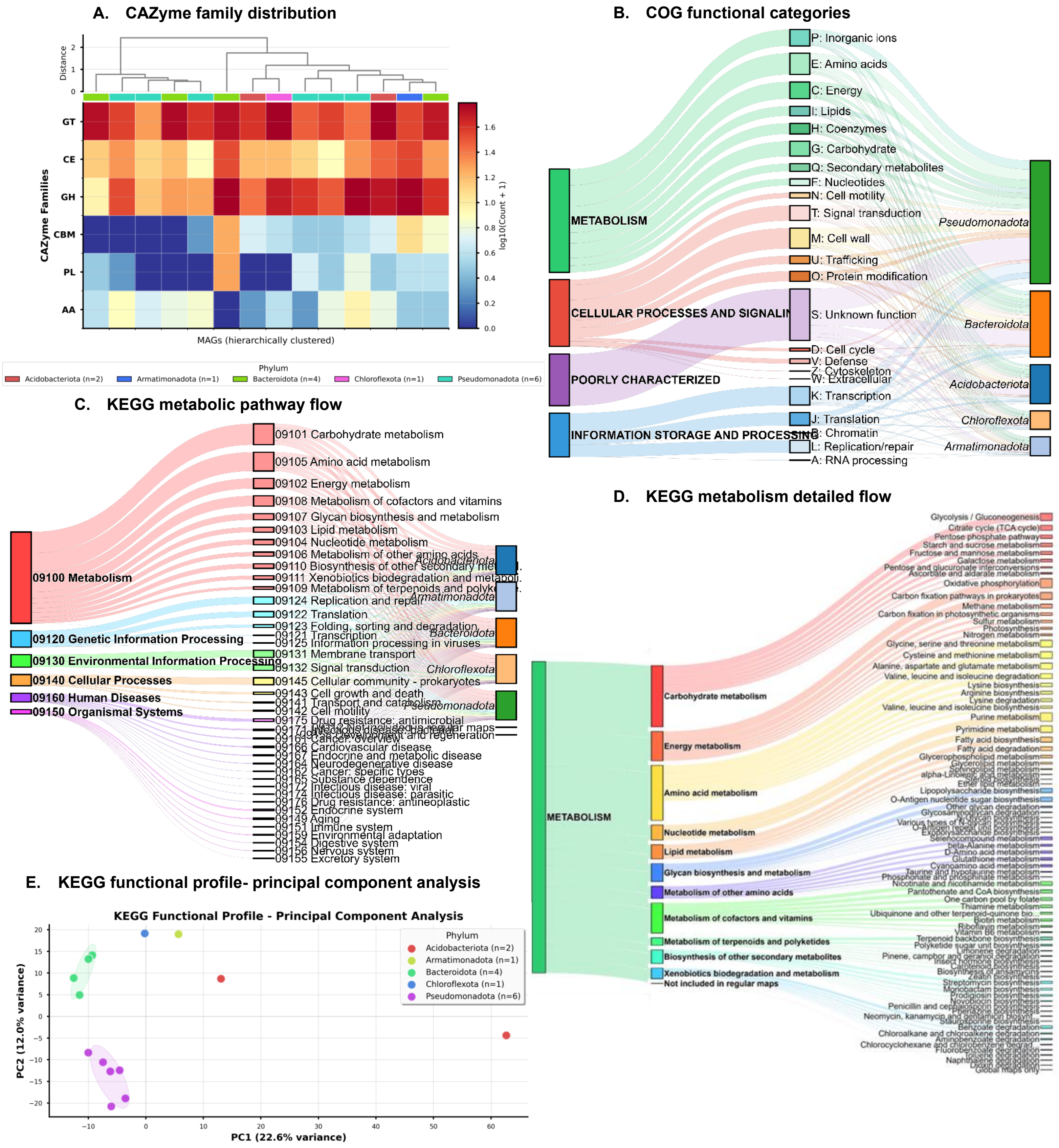


**Figure S6:** Functional characterization of novel MAGs from plant samples. **(A)** Distribution of carbohydrate-active enzyme (CAZyme) families across novel MAGs, clustered by genome and colored by phylum. **(B)** COG functional category assignments for predicted proteins, with the percentage of functions related to metabolism, cellular processes, information storage, and poorly characterized proteins. **(C)** KEGG metabolic pathway flow, connecting high-level functional categories such as metabolism, cellular processes and others to the taxonomic groups in which they are found. **(D)** Breakdown of individual pathways within the "Metabolism" category. **(E)** Principal component analysis (PCA) of KEGG functional profiles, with clustering of MAGs by phylum.

### Figure S7

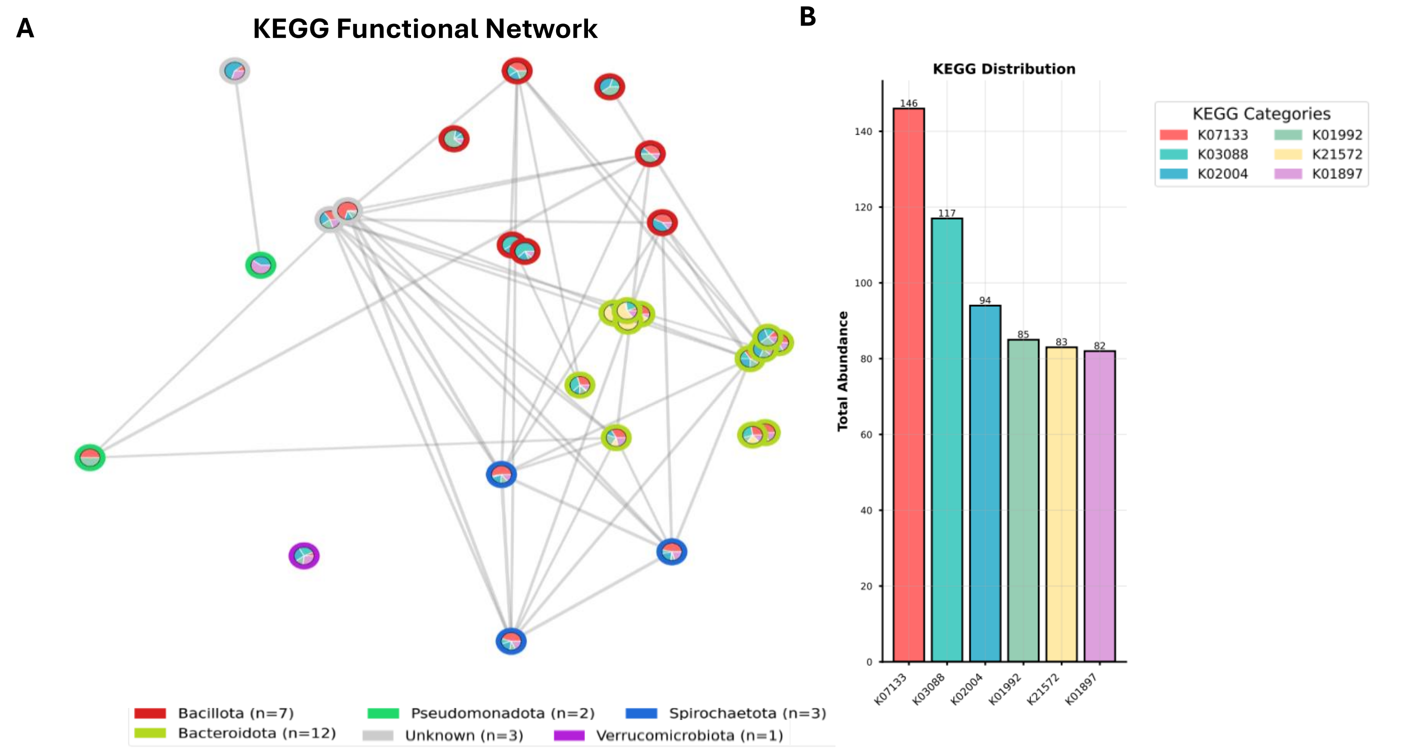


**Figure S7:** KEGG functional network: Panel A shows the KEGG pathway network where each node is MAG, colored by phylum. The connections between nodes indicate shared metabolic pathways - the more pathways two organisms share, the stronger their connection. Each node also contains a pie chart showing the KEGG pathway distribution for that MAG. Panel B summarizes the overall KEGG pathway distribution across all MAGs, corresponding to the categories shown in the node pie charts.

### Figure S8

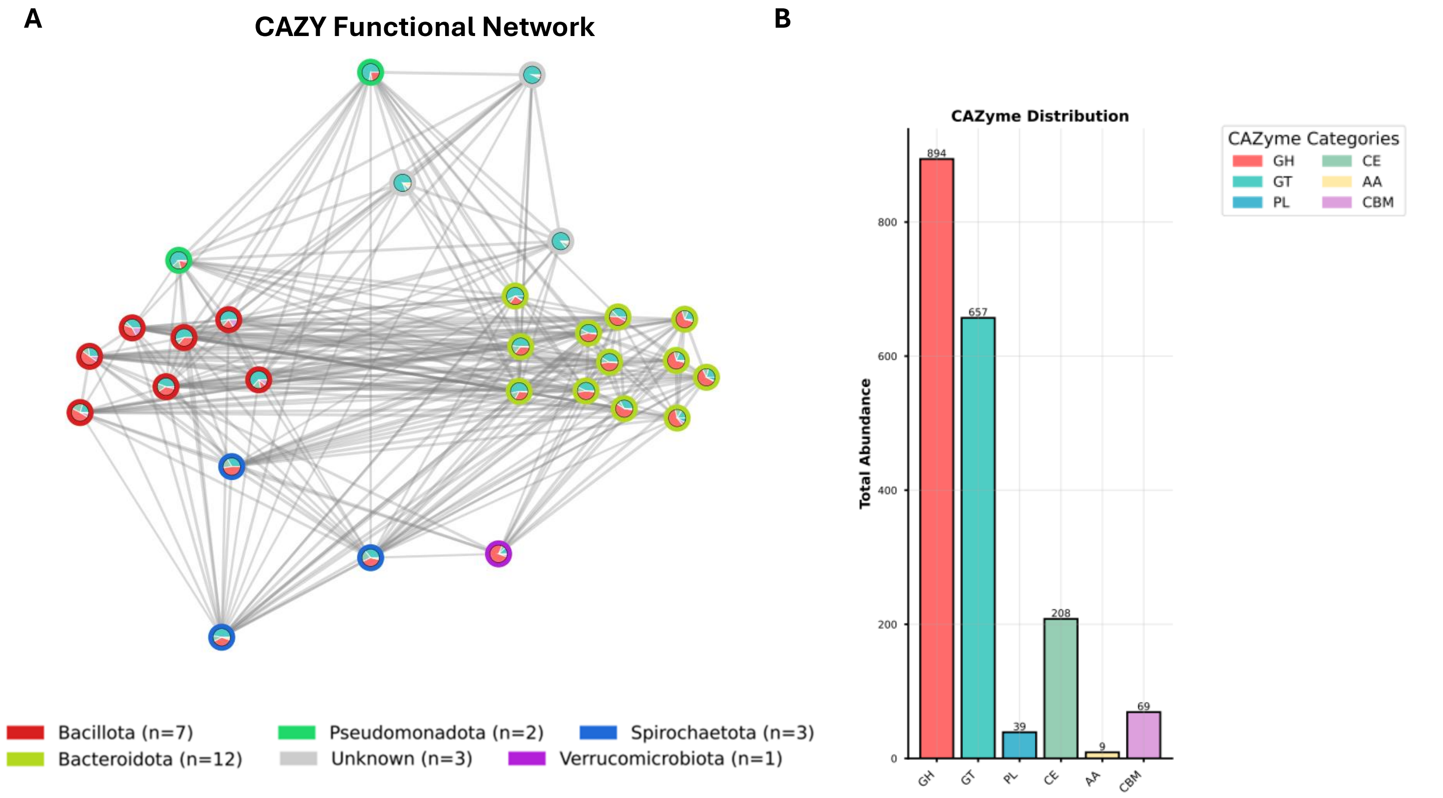


**Figure S8:** CAZYme functional network. (A) CAZYme functional network where every node represents a MAG colored according to phylum, with connections between nodes indicating shared CAZyme families (stronger connections indicate more shared families), and each node containing a pie chart displaying the CAZyme distribution for that MAG. (B) Summary of the general distribution of CAZyme categories across all MAGs, corresponding to the node pie chart categories shown in panel A.

### Figure S9

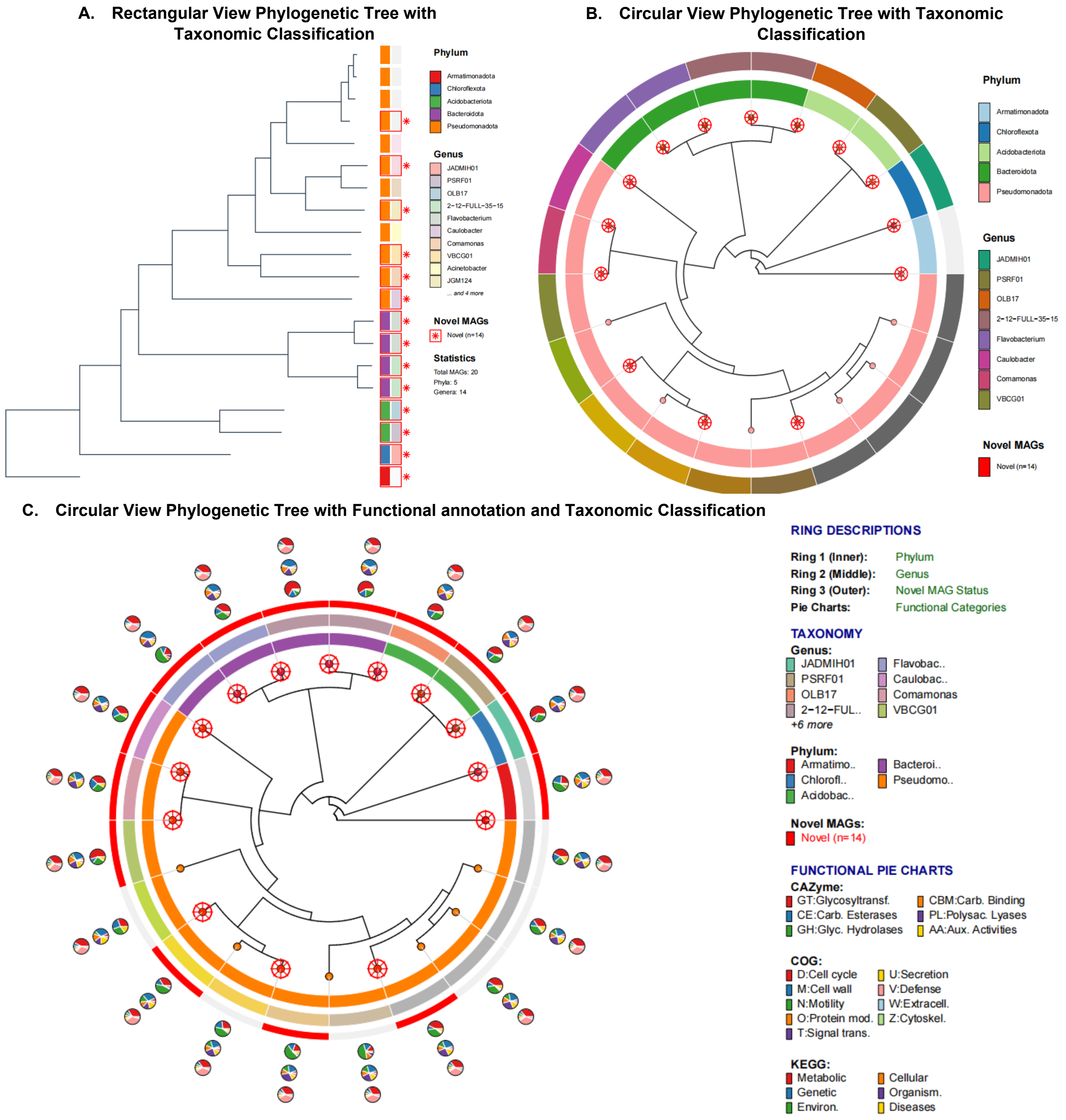


**Figure S9:** Phylogenetic placement and taxonomic classification of novel MAGs from the plant-associated dataset. (A) A rectangular phylogenetic tree with taxonomic classification at the phylum and genus levels; new MAGs are marked in red. (B) A circular phylogenetic tree that shows taxonomic classification by phylum (the outer ring) and genus (the inner ring), with new MAGs marked in red.(C) A circular phylogenetic tree that combines functional annotation with taxonomic classification. The circular phylogenetic tree shows different annotation layers, from the inner ring to the outer ring. The phylum classification for each MAG is shown in Ring 1 (the innermost ring). The second ring (middle) shows the genus classification. Ring 3 (the outer solid bars) indicates whether the MAG is new, with red bars highlighting newly discovered genomes. Three concentric functional annotation layers are displayed outside of these rings. The innermost functional ring shows CAZyme classes, the middle functional ring shows COG functional categories, and the outermost functional ring shows KEGG pathway categories. The pie charts around these functional categories show how they relate to each MAG.

#### Figure S10

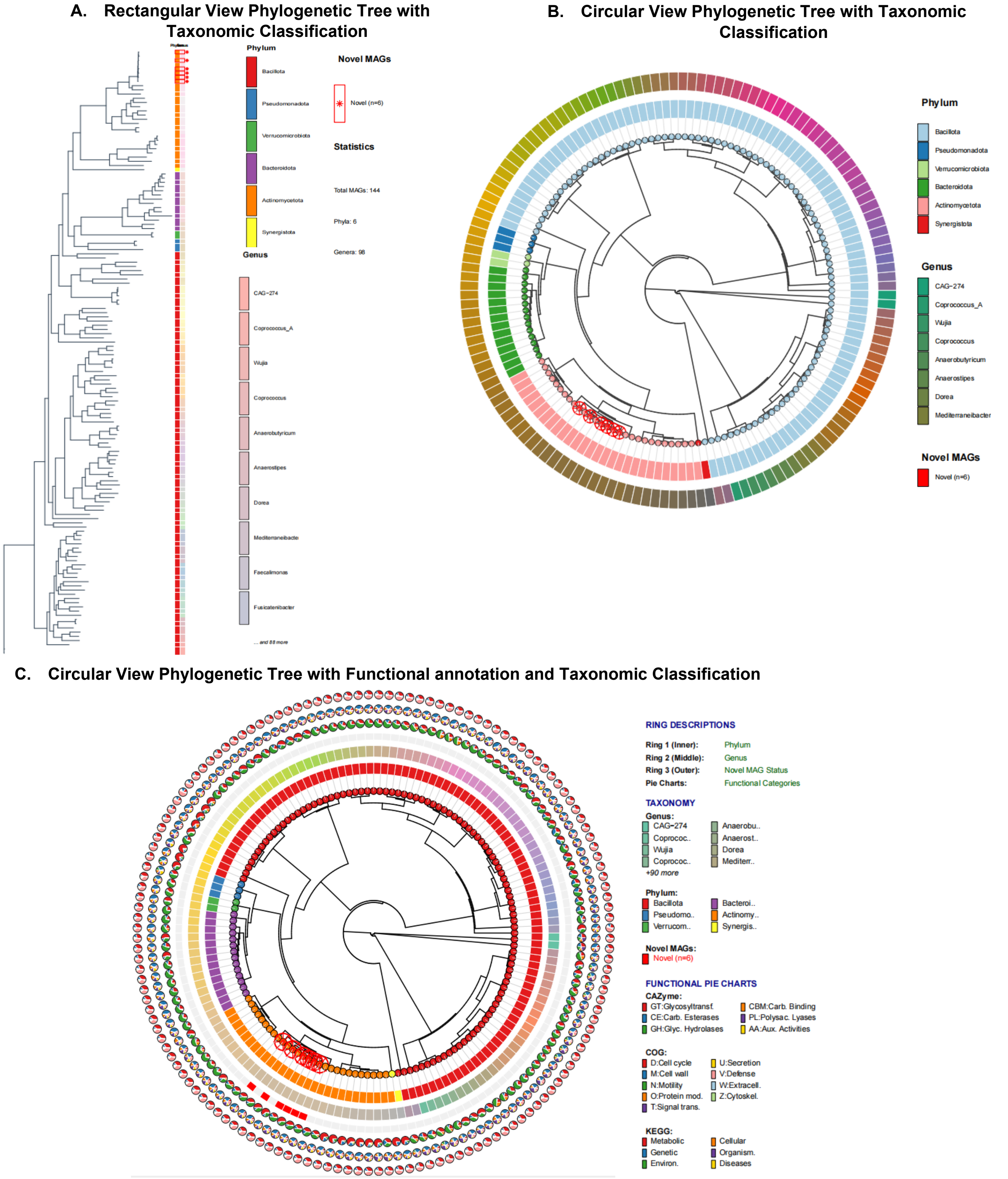


**Figure S10:** Phylogenetic placement and taxonomic classification of novel MAGs from the human-associated dataset. (A) A rectangular phylogenetic tree with taxonomic classification at the phylum and genus levels; new MAGs are marked in red. (B) A circular phylogenetic tree that shows taxonomic classification by phylum (the outer ring) and genus (the inner ring), with new MAGs marked in red. (C) A circular phylogenetic tree that combines functional annotation with taxonomic classification. The circular phylogenetic tree shows different annotation layers, from the inner ring to the outer ring. The phylum classification for each MAG is shown in Ring 1 (the innermost ring). The second ring (middle) shows the genus classification. Ring 3 (the outer solid bars) indicates whether the MAG is new, with red bars highlighting newly discovered genomes. Three concentric functional annotation layers are displayed outside of these rings. The innermost functional ring shows CAZyme classes, the middle functional ring shows COG functional categories, and the outermost functional ring shows KEGG pathway categories. The pie charts around these functional categories show how they relate to each MAG.

#### Figure S11

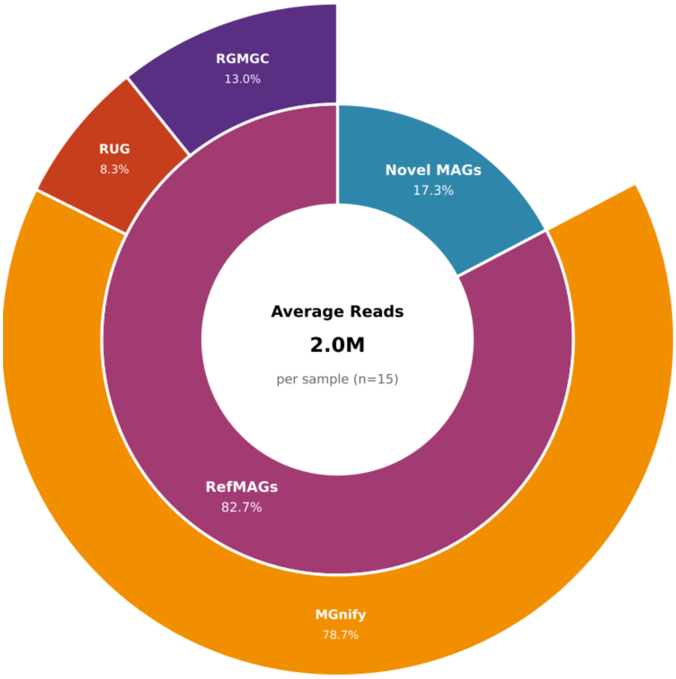


**Figure S11:** Hierarchical distribution of sequencing reads assigned to metagenome-assembled genomes (MAGs) following Kraken2 database enhancement. The inner ring displays the primary classification between novel MAGs identified in this study (n=28) and reference MAGs from published rumen microbiome studies (n=908). The outer ring shows the distribution of RefMAG-assigned reads across three major databases: MGnify (European Bioinformatics Institute), RUG (Rumen and Upper Gastrointestinal tract), and RGMGC (Rumen Gene Catalog). Values represent the average of reads across all 15 samples analyzed. RefMAGs contributed more classified reads than novel MAGs, with MGnify representing the dominant source of reference genomes.
